# Molecular characterization of Cdh12-SCON conditional knockout mice reveals unexpected splicing changes

**DOI:** 10.64898/2025.12.09.693148

**Authors:** Mayke A.C. ten Hoor, Margot M. Linssen, Conny M. Brouwers, Jill W.C. Claassens, Jaap Mulder, Peter Hohenstein

## Abstract

Functional validation of candidate genes in congenital anomalies of the kidneys and urinary tract (CAKUT) and other disorders is essential for translating genetic discoveries into clinical applications. Conditional knockout mouse models are indispensable for studying gene function in complex organ systems. The Short Conditional intrON (SCON) system accelerates the generation of such models by inserting the artificial SCON into a coding exon. SCON is designed to splice out after transcription, without affecting gene expression. Upon Cre activity, SCON is converted into the ΔSCON allele which cannot be spliced out, introducing premature termination codons (PTCs) to inactivate the gene. Previous validation of the SCON system in mice has focused primarily on phenotypic outcomes. Here, we provide a molecular characterization of the SCON system in Cdh12 – a candidate gene implicated in kidney damage in CAKUT. We found that both Cdh12^SCON^ and Cdh12^ΔSCON^ alleles caused unintended skipping of the exon downstream of the insertion site, culminating in a frameshift and PTC. Consequently, the Cdh12^SCON^ allele led to a ~25% reduction in mRNA expression, indicating that it was not transcriptionally inert as designed. Despite unintended exon skipping, the Cdh12^ΔSCON^ allele still effectively suppressed mRNA expression. These findings reveal previously unrecognized splicing artifacts of the SCON system and underscore the need for transcript-level characterization before utilizing artificial intron-based conditional alleles for functional studies.

## 1. Introduction

Congenital anomalies of the kidneys and urinary tract (CAKUT) comprise a heterogenous group of anomalies that vary significantly in severity, presentation, and outcome. Although advances in clinical care and diagnostics have improved prognosis, most cases remain genetically unexplained. Multiple large-scale initiatives are actively working to identify and prioritize candidate genes potentially involved in CAKUT and its disease progression (Vendrig et al., 2025). However, in vivo validation of these candidates is pursued to establish true causality and functional relevance to support their use in clinical practice (Richards et al., 2015).

Genetically engineered mouse models remain indispensable tools for studying gene function, particularly in the context of examining diseases whose pathogenesis arises from coordinated interactions between multiple cell types and organ systems. The pathogenesis of CAKUT relies on two main processes: the extent of initial organ maldevelopment and the subsequent progression of secondary kidney damage. Both processes are subject to complex spatiotemporal regulation at the genetic level (Jain & Chen, 2019). Therefore, conditional knockout approaches, which allow for gene inactivation in specific tissues or developmental stages, are particularly valuable in CAKUT research (Vendrig et al., 2025). The Cre-Lox recombination system is widely used to achieve such conditional control of gene expression, enabling researchers to dissect gene mechanisms in great detail.

To streamline the generation of conditional knockout alleles, the Short Conditional intrON (SCON) system was previously developed (Wu et al., 2022). SCON is a CRISPR/Cas9-based method that introduces a 189 bp artificial intron into the coding region of a target gene and is as efficient as CRISPR/Cas9-mediated gene tagging. From 5′ to 3′, the SCON cassette is composed of a splice acceptor (SA), three termination codons, the first loxP site, a branch point (BP), the second loxP site, a polypyrimidine tract, and a splice donor (SD). These elements should enable splicing out of SCON after transcription without affecting native mRNA expression. Upon Cre-mediated recombination, the BP is excised and a residual 55 bp sequence remains, hereafter called ΔSCON. The BP excision should impede splicing out of ΔSCON, disrupting mRNA translation to protein due to the introduction of premature termination codons (PTCs) in all three open reading frames and effectively silencing the gene (Wu et al., 2022).

The functionality of the SCON system comes with a challenge; it introduces artificial splice sites that could interfere with the complex process of pre-mRNA splicing. This process depends on the precise recognition of SD and SA sites, BPs, and other regulatory elements in the surrounding genomic architecture (Shenasa & Hertel, 2019). Since SCON introduces these elements, it is essential to characterize the resulting mutant alleles to validate their functionality and ensure that the intended disruption occurs without generating unintended splicing events. Despite its promise, the evaluation of SCON has been predominantly conducted at the phenotypic level, without detailed transcript-level analysis (Wu et al., 2022). Hence, the molecular consequences of SCON, particularly in multi-exon genes, remained unknown.

To evaluate the application of the SCON system and use it in studying CAKUT, we targeted the Cdh12 gene (Wu et al., 2022). CDH12 was previously implicated as a candidate modifier gene influencing the extent of kidney damage in a subset of CAKUT patients (van der Zanden et al., 2021). Cdh12 knockout mice generated by the Knockout Mouse Project (MGI:6293911) showed only mild phenotypes, limited to coat abnormalities in 3 out of 7 males. This minimal phenotype observed emphasizes that phenotypic assessment alone may overlook unexpected molecular disturbances, positioning Cdh12 as an ideal gene for uncovering unintended splicing effects. In this study, we provide a comprehensive molecular characterization of Cdh12 transcripts in Cdh12 mutants carrying the SCON or ΔSCON cassette, using RT-PCR and RT-qPCR. Our findings provide insight into the behavior of SCON in a multi-exon gene and highlight key considerations for its application in future genetic studies.

## 2. Materials and methods

### 2.1 Mouse strains and ethical compliance

All mice were housed at 20–22°C in individually ventilated cages, under a 12-h light/dark cycle with ad libitum access to water and food (standard RM3 chow; SDS, Essex, United Kingdom). The animal care and experimental procedures were approved by the Animal Welfare Body Leiden and are in accordance with the Dutch Experiments on Animals Act and EU Directive 2010/63/EU. The mouse strains C57BL/6JLumc and EIIa-Cre (B6.Cg-Tg(EIIa-cre)C5379Lmgd/Jlumc; MGI:2137691) backcrossed >30 times to C57BL/6JLumc were maintained within our animal facility. The generation of the Cdh12^SCON^ (C57BL/6JLumc-Cdh12^em1Lumc^) was described previously (Wu et al., 2022) (Fig.1). The Cdh12^ΔSCON^ (C57BL/6JLumc-Cdh12^em1.1Lumc^) was generated by crossing Cdh12^SCON/+^ with EIIa-Cre mice that carry a Cre transgene under the control of the adenovirus EIIa promoter. This promotor is active in preimplantation mouse embryos in a mosaic pattern, thus generating mosaic Cdh12^ΔSCON^ mice. The mosaic Cdh12^ΔSCON^ mice were used for further breeding to obtain germline transmission, which was confirmed by the presence of ΔSCON and the absence of the Cre allele.

### 2.2 Genotype assessment and sequencing

Genotypes were determined by PCR using genomic DNA extracted from ear clips. Biopsies were digested overnight at 55°C in lysis buffer (50 mM Tris pH 8.0, 100 mM EDTA, 100 mM NaCl, 1% SDS, 0.5 mg/mL Proteinase K). Genomic DNA was precipitated with saturated NaCl and isopropanol, washed with 70% ethanol, and rehydrated in TE^-4^ buffer (10 mM Tris-HCl, 0.1 mM EDTA, pH 8.0). Primers were designed to flank the insertion site, enabling the discrimination of alleles based on amplicon size: 354 bp for the wildtype, 543 bp for the Cdh12^SCON^, and 409 bp for the Cdh12^ΔSCON^ allele (Fig. 1B). For confirmation of loss of Cre in constitutive Cdh12^ΔSCON^ mice, primers in the Cre construct were used yielding a product for Cre presence and no product in case of absence. PCR amplification was carried out using Dreamtaq DNA polymerase (#EP0713, ThermoFisher). For Sanger sequencing, PCR products were purified from agarose gel. The primers and thermocycling conditions are presented in Table S1.

**Fig. 1.**
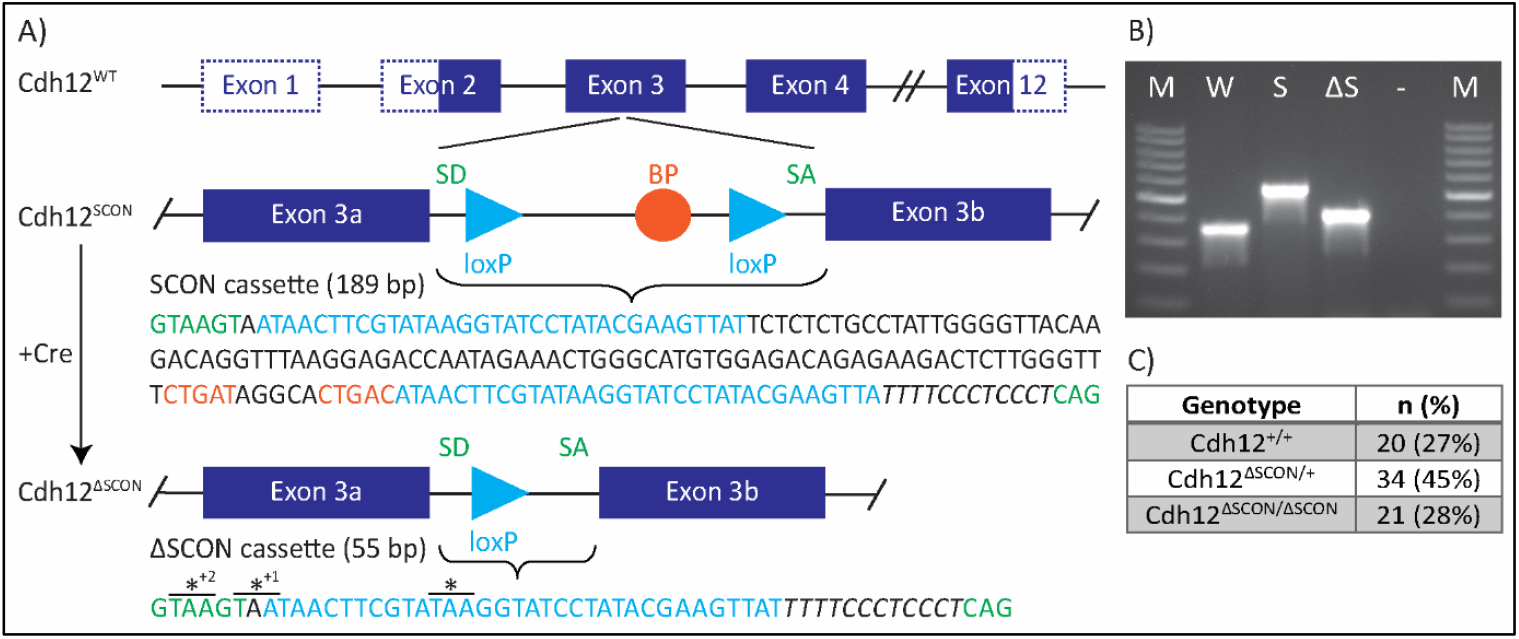
Generation of Cdh12^SCON^ and Cdh12^ΔSCON^ mice. A) Transgene structure of Cdh12 exon 3. The exons in blue are translated and the dotted lines indicate untranslated regions. Nucleotide colors correspond to functional elements in the sequence and is annotated as follows: the 189 bp SCON cassette contains a splice donor (SD; green), splice acceptor (SA; green), branch point (BP; orange), polypyrimidine tract (italic), and flanking loxP sites (blue). After Cre recombination, the 55 bp ΔSCON remnant retains the SD (green), SA (green), one loxP site (blue), and the polypyrimidine tract (italic), introducing premature termination codons (*) in multiple reading frames. B) Genotype identification of Cdh12 gene-edited mice. Lane annotations: M: 100 bp marker; W: Cdh12^+/+^; S: Cdh12^SCON/SCON^; ΔS: Cdh12^ΔSCON/ΔSCON^. C) Mendelian ratios from crosses of Cdh12^ΔSCON/+^ x Cdh12^ΔSCON/+^

### 2.3 RNA isolation

Embryos were generated by timed, overnight matings of either Cdh12^SCON/+^ x Cdh12^SCON/+^ or Cdh12^ΔSCON/+^ x Cdh12^ΔSCON/+^ animals. Pregnant females were sacrificed on the morning of E16.5 by cervical dislocation to collect the embryos. The embryos were removed from their amnion sac, culled, and stored in ice cold phosphate-buffered saline until their forebrain (excluding the olfactory bulb) was dissected (Fig. 2A). Total RNA was isolated and purified from the forebrain using the RNeasy Plus Mini Kit (Qiagen; 74134). RNA concentration was measured with NanoDrop and 500 ng of total RNA was reverse transcribed into cDNA using NEB’s standard first strand synthesis protocol with M-MulV reverse transcriptase (NEB; M0253), 60 µM random primer mix (NEB; S1330), 10 mM dNTP, and RNase inhibitor (NEB; M0314).

**Fig. 2.**
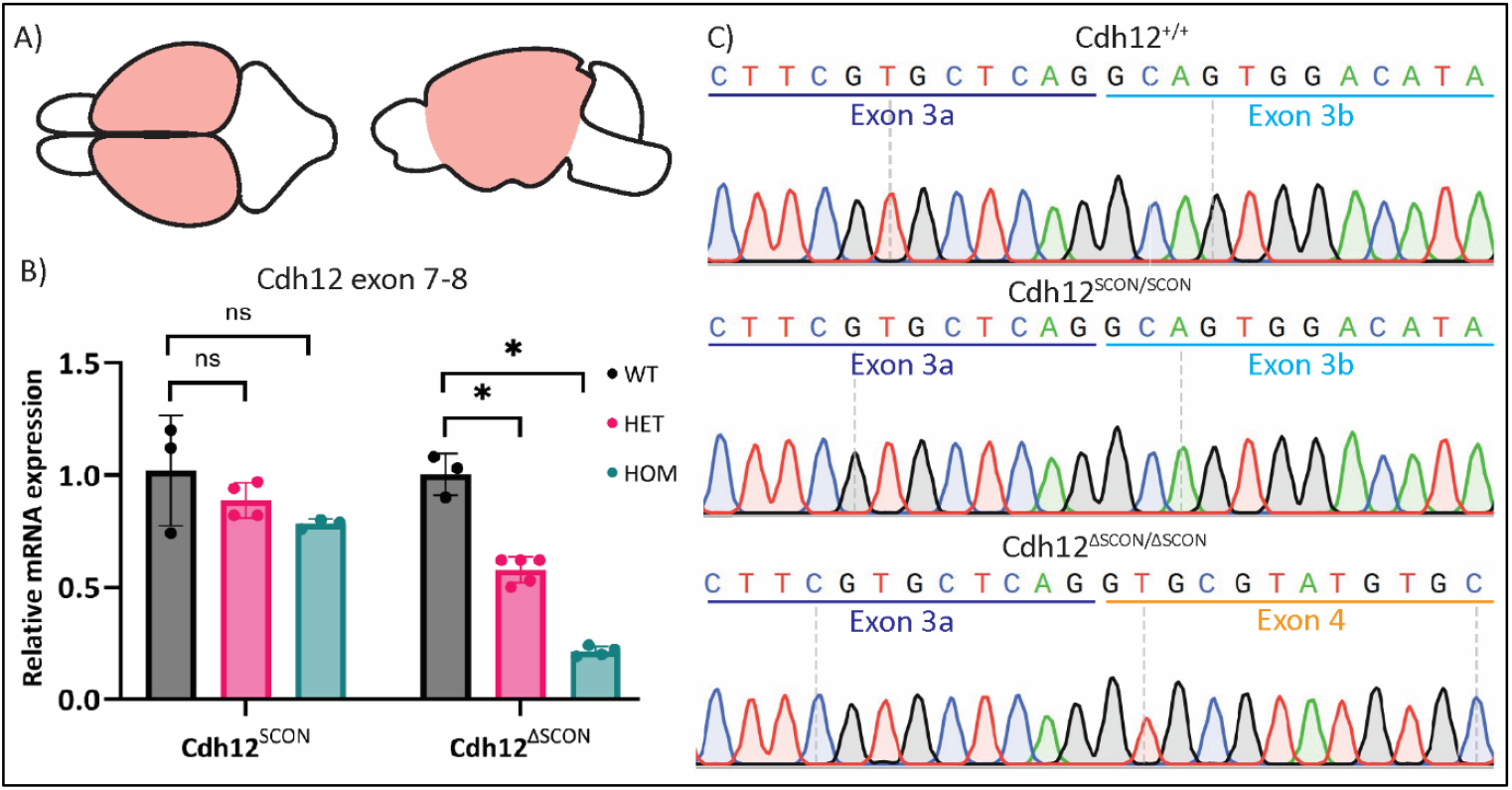
Cdh12 expression relative to Actb in the E16.5 embryonic forebrain. A) top and side view of the embryonic brain, the RNA was isolated from the E16.5 embryonic forebrain region highlighted in pink. B) Relative mRNA expression of Cdh12 for assays targeting the exon 7-8 junction for Cdh12^SCON^ and Cdh12^ΔSCON^ embryos. C) Sanger sequencing revealed normal mRNA splicing of exon 3 in Cdh12^+/+^ and Cdh12^SCON/SCON^. In contrast, exon 3b skipping was observed in Cdh12^ΔSCON/ΔSCON^. *p<0.001, one-way ANOVA. Cdh12^SCON^: WT (n=3), HET (n=4), HOM (n=3); Cdh12^ΔSCON^: WT (n=3), HET (n=5), HOM (n=4). Data are presented as the mean ± SEM

### 2.4 RT-qPCR

The qPCR mix was prepared using PrimeTime™ Gene Expression Master Mix (Integrated DNA Technologies; 1055772) with a final concentration of 500 nM primers and 250 nM probe from the predesigned PrimeTime™ qPCR probe assays for Cdh12 exon 7-exon 8 (Integrated DNA Technologies; Mm.PT.58.10546922) or Cdh12 exon 3-exon 4 (Integrated DNA Technologies; Mm.PT.58.42326005) with reference gene Actb (Integrated DNA Technologies; Mm.PT.39a.22214843.g) in each reaction. qPCR was performed in triplicate per biological sample using 0.5 ng cDNA input. Quantitation of Cdh12 was performed using the LightCycler® 480 Instrument II (Roche; 05015243001) with the following thermal cycling conditions: initial denaturation cycle at 95 °C for 3 min, and 45 cycles at 95 °C for 15 s and 60 °C for 1 min. The relative gene expression levels were calculated with respect to the Cdh12^+/+^ group, employing the 2^−ΔΔCT^ method.

### 2.5 RT-PCR

To analyze the nucleotide sequence of the Cdh12^SCON^ and Cdh12^ΔSCON^ mRNA, an initial RT-PCR was performed using primers spanning from exon 2 to exon 12. Amplification was carried out using Q5 High-Fidelity DNA Polymerase (NEB; M0191) and products were purified from agarose gel for Sanger sequencing. For more detailed isoforms characterization, two RT-PCR panels were designed. The first panel consisted of exon-exon spanning assays that surrounded the SCON insertion site, and the second panel entailed assays with one primer in the SCON/ΔSCON region and the other in upstream or downstream exons. These assays were performed on cDNA from Cdh12^+/+^, Cdh12^SCON/SCON^, and Cdh12^ΔSCON/ΔSCON^ E16.5 embryonic forebrains. A synthetic gene fragment (gBlock; Integrated DNA technologies) containing exons 2-7 with the ΔSCON cassette in exon 3 served as a positive control (supplementary information). MilliQ water was used as a no-template negative control. Each reaction contained 25 ng cDNA or 10,000 synthetic gene fragment copies. Primer sequences and PCR cycling conditions are listed in Table S1.

### 2.6 Splice site prediction

Splice site prediction was carried out using Spliceator v2.1(Scalzitti et al., 2021) and Human Splicing Finder (Desmet et al., 2009) to evaluate the effect of SCON or ΔSCON insertion in exon 3 on native SD and SA sites. The input sequence consisted of exon 3, exon3a-SCON-exon3b, or exon3a-ΔSCON-exon3b with 600 bp of flanking intronic sequence on both sides for the Cdh12^WT^, Cdh12^SCON^, or Cdh12^ΔSCON^ allele, respectively. For Spliceator, the 600 model was applied and the corresponding reliability scores were recorded. From the Human Splicing Finder, both the HSF matrix and MaxEnt scores were extracted.

### 2.7 Statistical analysis

Data are presented as mean ± SEM. Normality was tested using the Shapiro–Wilk test. Groups were compared using the one-way ANOVA followed by Tukey’s post hoc test. A value of p < 0.05 was considered statistically significant. All experiments were carried out by personnel blinded to genotype. The experimental unit was defined as a single embryo.

## 3. Results

### 3.1 Generation and identification of Cdh12^SCON^ and Cdh12^ΔSCON^ mice

The murine Cdh12 gene spans ~478 kb on chromosome 15. It comprises 12 exons, with the ATG start codon located in exon 2 and the TGA termination codon located in exon 12. To construct the Cdh12^SCON^ allele, SCON was inserted 160 bp downstream of the start of exon 3, as described previously (Wu et al., 2022) (Fig. 1A). Exon 3 was selected as it was previously the target of Cdh12 knockout mice (MGI:6293911), is common to all known protein coding isoforms, and meets the three conditions of a SCON targetable site: 1) exon positioned within the first 50% of the protein-coding sequence; 2) contains a stringent (MAGR, (A/C)-A-G-(A/G)) or flexible (VDGN, (A/G/C)-(A/T/G)-G-(A/T/G/C)) splice junction consensus sequence; 3) exon has a length of >120 bp with 5′ and 3′ split exons >60 bp (Wu et al., 2022). The Cdh12^SCON/+^ mice did not exhibit a phenotype and were able to reproduce. Hereafter, the parts of exon 3 upstream and downstream of the insertion site are referred to as exon 3a and exon 3b, respectively.

To verify that the floxed sequence in the SCON cassette could be recombined by Cre in vivo, Cdh12^SCON/+^ mice were crossed with EIIa-Cre mice, which express Cre in nearly all tissues of pre-implantation mouse embryos in a mosaic pattern. These matings successfully generated mosaic Cdh12^ΔSCON^ mice. Constitutive Cdh12^ΔSCON^ mice were successfully generated by crossing the mosaic Cdh12^ΔSCON^ mice with wildtype mice (Fig. 1B). The resulting Cdh12^ΔSCON/+^ mice displayed no phenotype and were fertile. Interbreeding of these mice yielded offspring with the anticipated Mendelian ratios (Fig. 1C).

### 3.2 Initial assessment of Cdh12 mRNA transcript integrity

For the initial assessment of whether the insertion of SCON or ΔSCON disrupts the integrity of Cdh12 mRNA transcripts, we performed RT-qPCR and RT-PCR analyses on mRNA isolated from the E16.5 embryonic forebrain of wildtype, Cdh12^SCON^, and Cdh12^ΔSCON^ mice (Fig. 2A). The embryonic forebrain was selected because of its documented expression of Cdh12 at this developmental stage (Mayer et al., 2010). RT-qPCR targeted the exon 7-8 junction to determine whether the insertion into exon 3 affects the continuity of downstream exons. To further evaluate full-length transcript integrity, RT-PCR was conducted using primers spanning exon 2 to exon 12, followed by Sanger sequencing of the amplified products. If the SCON insertion is functionally silent at the transcript level, it should neither disrupt normal mRNA expression nor be present in the mature mRNA. Consistent with this expectation, RT-qPCR analysis revealed no significant difference in Cdh12 exon 7-8 junction expression between wildtype, heterozyous, and homozygous Cdh12^SCON^ embryos (p=0.192) (Fig. 2B). In addition, the sequencing results showed that the full-length mRNA sequence of Cdh12^SCON/SCON^ is identical to the sequence identified in Cdh12^+/+^(Fig.2C). These findings indicate that SCON insertion does not impair downstream transript levels and appears to be succesfully spliced from the mRNA.

The ΔSCON insertion was expected to disrupt full length Cdh12 mRNA expression due to the introduction of PTCs, which would be retained in the mature transcript. In alignment with this expectation, RT-qPCR analysis showed a significant reduction in Cdh12 mRNA levels at the 7-8 junction: expression dropped to 57.80% ± 0.03% (p<0.001) in Cdh12^ΔSCON/+^ and to 21.25% ± 0.01% (p<0.001) in Cdh12^ΔSCON/ΔSCON^ embryos compared to Cdh12^+/+^ controls (Fig. 2B). However, further transcript analysis uncovered an unintended splicing event in Cdh12^ΔSCON/ΔSCON^ mice (Fig. 2C). This event involved the splicing out of the ΔSCON cassette along with the entire 136 bp of exon 3b, resulting in a new isoform that introduced a frameshift culminating in a PTC in exon 4. Thus, although the observed PTC arose from aberrant splicing rather than ΔSCON retention, the insertion nonetheless led to impaired Cdh12 mRNA integrity downstream of the target site as intended.

### 3.3 mRNA isoform analysis

The identification of an unintended splicing event in the Cdh12^ΔSCON/ΔSCON^ mRNA prompted us to conduct an in-depth analysis of Cdh12 mRNA isoforms in both Cdh12^ΔSCON/ΔSCON^ and Cdh12^SCON/SCON^ mice. To characterize Cdh12 transcripts with high sensitivity, we performed two RT-PCR panels. The first panel entailed exon-exon spanning assays surrounding the SCON insertion site (Fig 3A, I-VII), and the second panel consisted of assays with one primer in SCON/ΔSCON region and the other in adjacent exons (Fig 3A, VIII-XIV). A synthetic gene fragment containing exons 2-7 with the ΔSCON cassette served as a positive control. Table 1 shows the predicted amplicon sizes based on the theoretical function of SCON and ΔSCON, as well as the observed amplicon sizes.

**Table 1.**
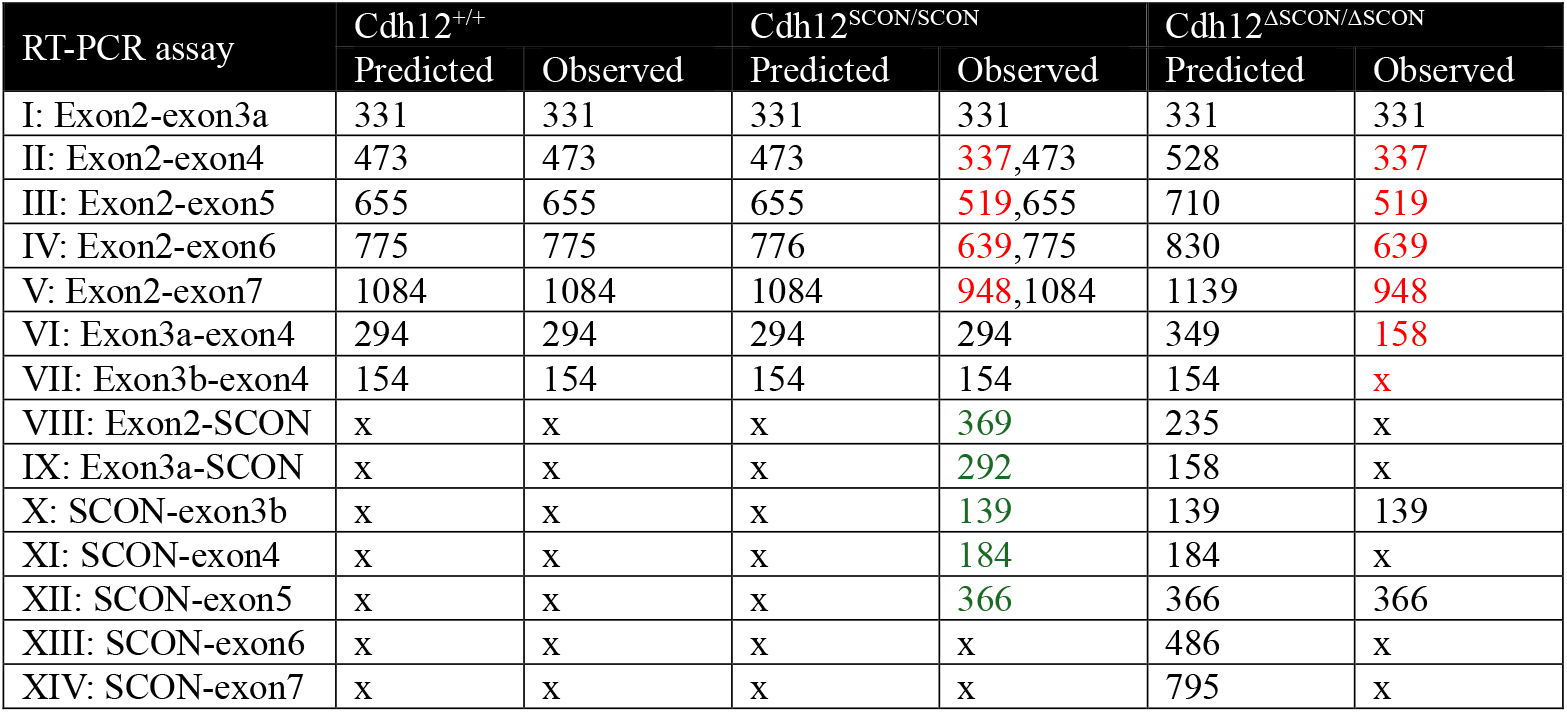
Predicted and observed amplicon length for all RT-PCR assays. Green and red text indicate SCON retention and exon skipping, respectively. Amplicons identified as intron retention are not included in the table

| RT-PCR assay | *Cdh12*<sup>+/+</sup> |  | *Cdh12*<sup>SCON/SCON</sup> |  | *Cdh12*<sup>ΔSCON/ΔSCON</sup> |  |
| --- | --- | --- | --- | --- | --- | --- |
|  | Predicted | Observed | Predicted | Observed | Predicted | Observed |
| I: Exon2-exon3a | 331 | 331 | 331 | 331 | 331 | 331 |
| II: Exon2-exon4 | 473 | 473 | 473 | 337,473 | 528 | 337 |
| III: Exon2-exon5 | 655 | 655 | 655 | 519,655 | 710 | 519 |
| IV: Exon2-exon6 | 775 | 775 | 776 | 639,775 | 830 | 639 |
| V: Exon2-exon7 | 1084 | 1084 | 1084 | 948,1084 | 1139 | 948 |
| VI: Exon3a-exon4 | 294 | 294 | 294 | 294 | 349 | 158 |
| VII: Exon3b-exon4 | 154 | 154 | 154 | 154 | 154 | x |
| VIII: Exon2-SCON | x | x | x | 369 | 235 | x |
| IX: Exon3a-SCON | x | x | x | 292 | 158 | x |
| X: SCON-exon3b | x | x | x | 139 | 139 | 139 |
| XI: SCON-exon4 | x | x | x | 184 | 184 | x |
| XII: SCON-exon5 | x | x | x | 366 | 366 | 366 |
| XIII: SCON-exon6 | x | x | x | x | 486 | x |
| XIV: SCON-exon7 | x | x | x | x | 795 | x |

**Fig. 3.**
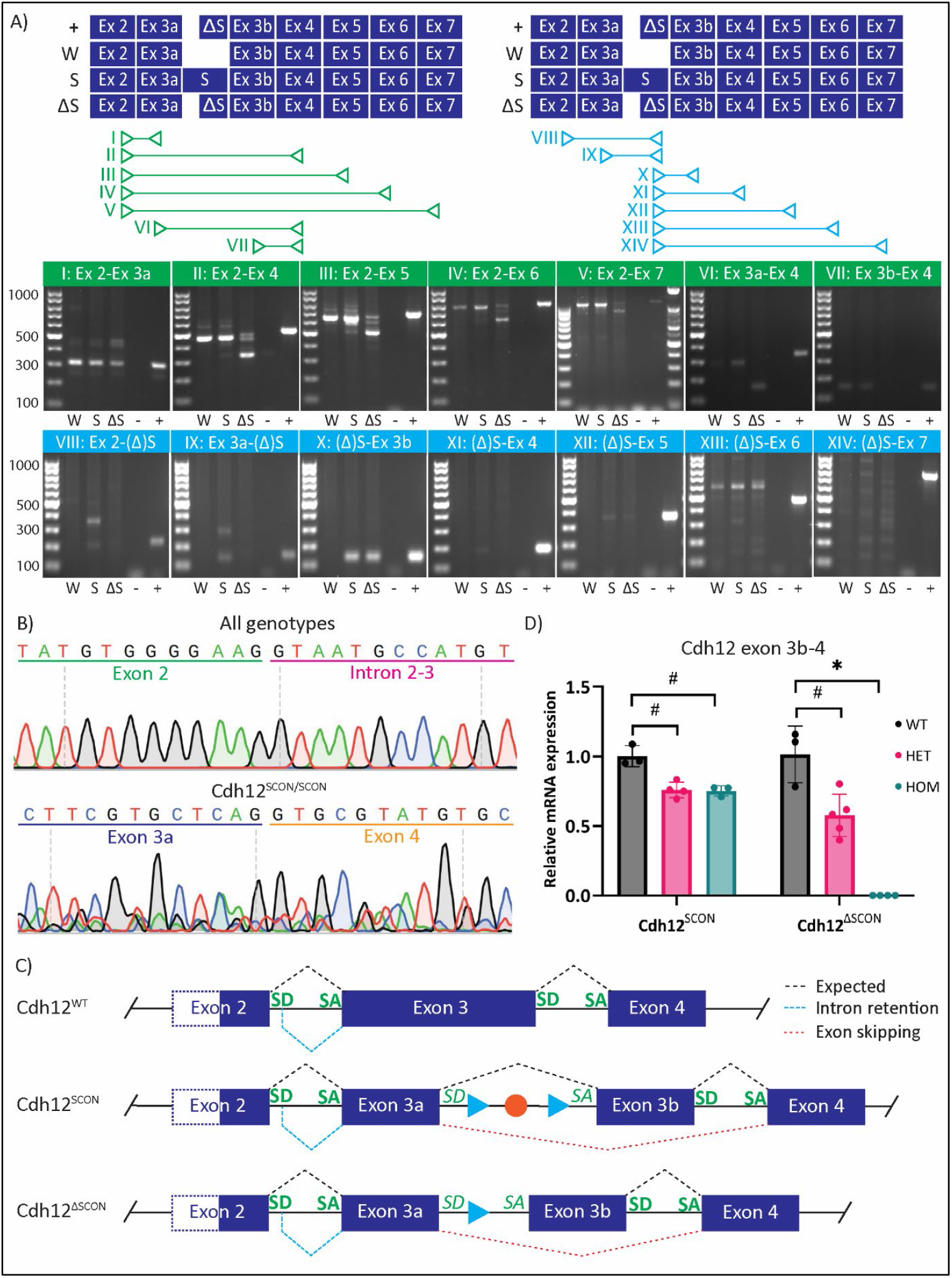
Isoform analysis of Cdh12^SCON/SCON^ and Cdh12^ΔSCON/ΔSCON^ mRNA. A) The RT-PCR assays for the isoform analysis depicted with the predicted mRNA sequence of the positive control (+), Cdh12^+/+^ (W), of Cdh12^SCON/SCON^ (S), Cdh12^ΔSCON/ΔSCON^ (ΔS) and the corresponding RT-PCR results on agarose gel. Lane annotations: W: Cdh12^+/+^; S: Cdh12^SCON/SCON^; ΔS: Cdh12^ΔSCON/ΔSCON^; − = MQ (negative control); + = synthetic gene fragment (positive control); Ladders: 100 bp marker used throughout, except for exon 2-exon 7 panel, which uses a 1 kb+ ladder on the right. B) Sanger sequencing revealed partial intron retention in all genotypes and exon 3b skipping in Cdh12^SCON/SCON^. C) Schematic illustration of identified isoforms. Native splice acceptors (SA) and splice donors (SD) in bold and artificial SA and SD in italic. D) Relative mRNA expression of Cdh12 for assays targeting the exon 3b-4 junction for Cdh12^SCON^ and Cdh12^ΔSCON^ embryos. #p<0.01, *p<0.001, one-way ANOVA. Cdh12^SCON^: WT (n=3), HET (n=4), HOM (n=3); Cdh12^ΔSCON^: WT (n=3), HET (n=5), HOM (n=4). Data are presented as the mean ± SEM

In RT-PCR assays flanking exon 2 and 3, bands above the expected amplicon length were detected in all genotypes (Fig. 3A, I-V). To confirm that these products were not related to the introduction of SCON or ΔSCON, these fragments were sequenced. The sequencing data revealed that these products are a previously undescribed isoform of Cdh12 (Fig. 3B, only first 12 bp shown). This isoform entailed the partial retention of 158 bp of intron 2-3 that leads to a PTC almost directly downstream of exon 2. The presence of this isoform in Cdh12^+/+^ implies that this isoform was unrelated to SCON or ΔSCON.

In Cdh12^SCON/SCON^ mice, the SCON cassette was expected to be spliced out, producing RT-PCR products identical to those of Cdh12^+/+^ controls. Consistent with this expectation, RT-PCR assays using primers flanking SCON showed that the majority of bands matched the Cdh12^+/+^ pattern (Fig. 3A, I-VII). However, an additional fragment of ~136 bp shorter was detected in Cdh12^SCON/SCON^ (Fig. 3A, II-V). Sanger sequencing revealed that this amplicon resulted from unintended skipping of both SCON and exon 3b, which is 136 bp in length (Fig. 3B). Furthermore, fragments were observed in most assays that included a primer within the SCON cassette, indicating low-level retention of SCON (Fig. 3A VIII-XII). Taken together, these results show that while SCON was correctly removed in most transcripts, some retained the cassette or exhibited aberrant splicing. All detected isoforms are summarized in Fig. 3C.

The ΔSCON cassette was expected to be retained in the mRNA of Cdh12^ΔSCON/ΔSCON^ mice, yielding amplicons of 55 bp longer than those from Cdh12^+/+^ controls and matching the positive control. Contrary to this, amplicons flanking ΔSCON-exon 3b were ~136 bp shorter than those from Cdh12^+/+^ controls, and no product was detected in the exon 3b-exon 4 assay (Fig 3A, I-VII). These results indicated that both ΔSCON and exon 3b were skipped during splicing. Consistent with this, no products were dectected in any assay that included a primer within ΔSCON – with two notable exceptions: the ΔSCON-exon 3b and ΔSCON-exon 5 assays (Fig 3A, X & XII). The presence of these two products contrast with the overall pattern of ΔSCON-exon 3b loss. The ΔSCON-exon 3b amplicon was confirmed to be mRNA-derived, as reverse transcriptase-negative controls showed no amplification, excluding genomic DNA contamination (Fig. S1). Although the ΔSCON-exon 5 fragment appeared at the predicted length – implying inclusion of exon 4 – the ΔSCON-exon 4 fragment was absent (Fig 3A, XI). Attempts to sequence the ΔSCON-exon 5 amplicon did not establish sufficient reads for analysis. Taken together, these findings confirm consistent ΔSCON-exon 3b skipping in Cdh12^ΔSCON/ΔSCON^ transcripts.

However, the origin of the ΔSCON-exon 3b and ΔSCON-exon 5 products, which deviate from this pattern, remains unresolved.

### 3.4 Quantification of exon 3b skipping

To quantifiy the extent of exon 3b skipping in Cdh12^SCON^ and Cdh12^ΔSCON^ mice, a RT-qPCR was conducted targetting the exon 3b-exon 4 junction. The qPCR analysis revealed a significant reduction in exon 3b-4 expression to 75.94% ± 0.03% (mean ± SEM; p=0.002) in Cdh12^SCON/+^ and to 75.27% ± 0.02% (mean ± SEM; p=0.003) in Cdh12^SCON/SCON^ compared to control levels (Fig. 3D). Expression in Cdh12^ΔSCON/+^ was decreased to 57.62% ± 0.07 (mean ± SEM; p=0.005) relative to controls, whereas no detectable expression was observed in Cdh12^ΔSCON/ΔSCON^ (p<0.001) (Fig. 3D). This analysis demonstrates that exon 3b was significantly lost in Cdh12^SCON/+^ Cdh12^SCON/SCON^ and Cdh12^ΔSCON/+^ mice, and was undetectable in Cdh12^ΔSCON/ΔSCON^ mice.

### 3.5 In silico splice score prediction

The observed skipping of exon 3b in a subset of Cdh12^SCON^ transcripts and all of Cdh12^ΔSCON^ transcripts suggests that splicing occurred from the SD within SCON or ΔSCON - hereinafter collectively referred to as (Δ)SCON - to the SA of exon 4, bypassing the native SD located downstream of exon 3b (Fig. 3C). To explore how splice site prediction tools would evaluate the splicing pattern, we conducted in silico analysesusing Spliceator and Human Splicing Finder, extracting splice site reliability scores from the former and HSF matrix and MaxEnt scores from the latter (Table 2). It is suggested that insertion of an artificial intron should not reduce native splice site scores by more than 1% (McBeath et al., 2023). If the occurrence of the unintended splicing event was predictable, we would expect the score for the native exon 3 SD to change by at least that threshold. However, the exon 3 SD score showed only a minor increase of 0.1% in splice site reliability, and the exon 3 SA score remained unchanged following (Δ)SCON insertion (Table 2). These findings indicate that the aberrant splicing event was not anticipated by the currently available prediction tools.

**Table 2.**
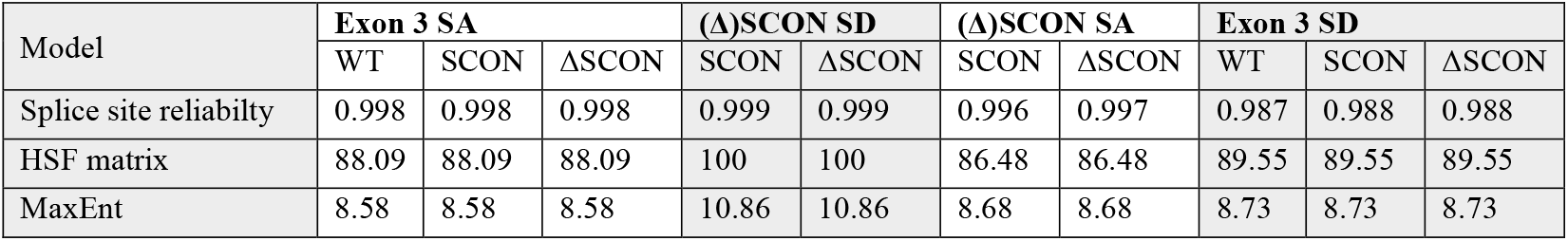
Spice site prediction scores.

| Model | Exon 3 SA |  |  | (Δ)SCON SD |  | (Δ)SCON SA |  | Exon 3 SD |  |  |
| --- | --- | --- | --- | --- | --- | --- | --- | --- | --- | --- |
|  | WT | SCON | ΔSCON | SCON | ΔSCON | SCON | ΔSCON | WT | SCON | ΔSCON |
| Splice site reliability | 0.998 | 0.998 | 0.998 | 0.999 | 0.999 | 0.996 | 0.997 | 0.987 | 0.988 | 0.988 |
| HSF matrix | 88.09 | 88.09 | 88.09 | 100 | 100 | 86.48 | 86.48 | 89.55 | 89.55 | 89.55 |
| MaxEnt | 8.58 | 8.58 | 8.58 | 10.86 | 10.86 | 8.68 | 8.68 | 8.73 | 8.73 | 8.73 |

Although the splicing pattern could not be predicted, it is noteworthy that the SD within (Δ)SCON consistently scored higher than the native exon 3 SD, suggesting that under certain conditions, the (Δ)SCON SD may be preferentially utilized over the native donor site.

## 4. Discussion

Genes implicated in disease through genomic studies often necessitate in vivo validation to support their use in clinical practice. In this study, the SCON system was employed to rapidly generate mice with a conditional knockout allele in the Cdh12 gene, which is a candidate gene for kidney damage in a subset of CAKUT patients. Functionally, the SCON allele should not interfere with protein expression, while after recombination the ΔSCON allele should interrupt functional protein expression. Although the original SCON study demonstrated the neutrality of SCON and hypomorphic effect of ΔSCON (Wu et al., 2022), it did not evaluate transcript-level consequences. Given that SCON introduces additional splice sites and Cdh12 knockout mice are not expected to display overt phenotypes, we decided to perform a detailed molecular characterization to validate SCON functionality. We conducted an analysis of Cdh12^SCON^ and Cdh12^ΔSCON^ mRNA using RT-PCR and RT-qPCR to assess the impact of these alleles on Cdh12 transcript processing.

Insertion of the 189 bp SCON cassette into exon 3 of Cdh12 successfully generated Cdh12^SCON^ mice (Wu et al., 2022). Although these mice did not display an overt phenotype and SCON was spliced out as intended in most transcripts, isoform analysis revealed occasional cassette retention or skipping of exon 3b. Both events introduce a PTC immediately downstream of exon 3a, consistent with the significant reduction of ~25% in RT-qPCR signal observed at the exon 3b-4 junction in both heterozygous and homozygous Cdh12^SCON^ mice. While protein-level effects could not be quantified due to the lack of antibody specificity, these transcript alterations suggest an undesirable hypomorphic impact of SCON on Cdh12 expression. By contrast, no significant reduction was observed at the exon 7-8 junction, underscoring the need for thorough transcript-level validation when assessing SCON alleles.

Following EIIa Cre-mediated recombination, Cdh12^ΔSCON^ mice were generated. No phenotype was observed in Cdh12^ΔSCON/ΔSCON^, although previously coat abnormalities were reported in a few male Cdh12 knockouts (MGI:6293911). However, the original description of this coat phenotype was unclear and consequently difficult to compare. As expected, the genomic presence of ΔSCON in heterozygous and homozygous Cdh12^ΔSCON^ mice exerted a hypomorphic effect on Cdh12 mRNA expression. Importantly, this reduction was not due to the intended ΔSCON retention but rather to the consistent skipping of both ΔSCON and exon 3b, which culminated in a PTC directly downstream of exon 3a. This position corresponds closely to where the PTC would occur if ΔSCON was retained. RT-qPCR revealed 21.25% residual Cdh12 transcript expression in Cdh12^ΔSCON/ΔSCON^ mice, but with complete exclusion of exon 3b. Although we could not perform protein-level validation due to the lack of antibody specificity, these findings suggest that ΔSCON exerts a biologically relevant hypomorphic effect at the transcript level, likely culminating in a Cdh12 null mutation.

ΔSCON was still detected as a separate entity with exon 3b, which is expected to be excised as an intron. Spliced introns can be rapidly degraded, processed into non-coding RNA intermediates, or stabilized as circular RNAs (Liao & Zheng, 2025). The presence of ΔSCON-exon 3b may reflect retention as non-coding or circular RNA but is unlikely to impede the generation of a Cdh12 null mutant.

Although the artificial intron target site in Cdh12 accords with recently established design guidelines (McBeath et al., 2023), we observed unintended skipping of (Δ)SCON-exon 3b. A similar phenomenon was reported with the DECAI artificial intron, where Cre recombination led to skipping of the exon downstream of the insertion site (Cassidy & Pelletier, 2023). These findings highlight the complexity of pre-mRNA splicing, which relies on precise splice site recognition by the spliceosome – a process influenced by multiple factors including the surrounding intron-exon architecture and splicing factors (Speakman & Gunaratne, 2024). Given that alternative SD selection is a significant driver of transcript diversification (Hicks et al., 2010), the strong (Δ)SCON SD can misguide the spliceosome, leading to preferential usage over the native exon 3 SD. However, in silico splice site predictions programs failed to anticipate the alternative SD selection. These results emphasize the importance of selecting a target site where unintended splicing would result in a frameshift and ideally introduce a PTC, minimizing the risk of functional protein production (McBeath et al., 2023).

The substantial exon 3b skipping observed in Cdh12^SCON^ mice differs from the initial study that demonstrated the neutrality of SCON when inserted into eGFP, Sox2, and Ctnnb1 (Wu et al., 2022). This discrepancy likely reflects the influence of genomic context. In the single-exon eGFP construct and Sox2 gene, the absence of competing splice sites simplifies splicing, whereas Ctnnb1 and Cdh12 are multi-exon genes with distinct intron-exon architectures. Moreover, the Ctnnb1^SCON^ allele is flanked by short introns (<250 bp), while the Cdh12^SCON^ allele is surrounded by introns that exceed 120 kb (Shenasa & Hertel, 2019). However, no quantitative mRNA or protein data were provided for Ctnnb1^SCON^, which prevents conclusions about unintended splicing events. Collectively, these findings underscore the critical role of genomic context in SCON behavior. Current prediction tools perform well for intronic variants but struggle with exonic changes (Smith & Kitzman, 2023), which is a significant limitation when working with artificial introns like SCON. More research is needed to understand how artificial introns affect the genomic context and to develop better tools for selecting effective target insertion sites.

In this study, RT-PCR isoform analysis was confined to exons 2-7. Splicing outside this region or isoforms involving intron retention within the long introns of the Cdh12 gene could therefore have been missed, particularly if these exceeded the size range amplifiable by PCR. While this analysis allowed for the identification of a previously undescribed isoform, long-read sequencing could provide a more complete view of the transcriptome.

Cdh12 is a single-pass transmembrane protein containing five extracellular cadherin (EC) repeats that mediate cell adhesion (Brasch et al., 2018). Despite extensive testing with multiple commercially available antibodies, consistent Cdh12 protein detection was unsuccessful, even in wildtype tissues. Given this limitation, we can only predict the potential impact of the PTC introduced by SCON retention or (Δ)SCON-exon 3b skipping on protein expression. This PTC likely triggers non-sense mediated mRNA decay, nascent protein degradation, or truncated protein production (Chemla et al., 2018), making full-length Cdh12 production improbable. The predicted truncated protein includes only the first 130 of 794 residues, covering most of the EC1 domain. While EC1 mediates dimerization via Trp2/Trp4 docking into a hydrophobic pocket of the partner EC1 domain (Brasch et al., 2018), the truncated Cdh12 lacks key hydrophobic pocket residues, rendering dominant-negative or gain-of-function activity highly unlikely (Tan et al., 2010). Nonetheless, residual or non-canonical activity cannot be ruled out without functional assays. Overall, these data suggest that SCON retention or (Δ)SCON-exon 3b skipping exerts a hypomorphic effect on Cdh12 protein expression.

In summary, this study presents a comprehensive molecular characterization of Cdh12^SCON^ and Cdh12^ΔSCON^ alleles, revealing that both led to unintended splicing events resulting in a frameshift and a PTC downstream of exon 3a. These findings demonstrate that SCON is not entirely neutral and ΔSCON could not be retained when inserted into the multi-exon Cdh12 gene. These splicing artifacts of the SCON system highlight the importance of characterizing transcript-level consequences before utilizing artificial intron-based conditional alleles for functional studies. Beyond validation of the model, analyzing mRNA isoform also deepens our understanding of how artificial introns behave in different genomic environments. As our understanding of pre-mRNA splicing advances and splice prediction tools continue to improve, the reliability and success of artificial intron technologies like SCON are expected to increase.

## Abbreviations

Actb: β-Actin
BP: Branchpoint
CAKUT: Congenital anomalies of the kidneys and urinary tract
Cdh12: Cadherin 12
Ctnnb1: Catenin β-1
SA: Splice acceptor
SCON: Short Conditional intrON
SD: Splice donor
Sox2: SRY-box transcription factor 2
PTC: Premature termination codon

## Acknowledgements

This study was funded by the Dutch Kidney Foundation through the ArtDECO consortium (no. 20OC002). J.M. is a member of the NL Research consortium Kidnie, which is supported by the Dutch Kidney Foundation, and of the European Reference Network for Rare Kidney Diseases (ERKNet).

## Author information

### Authors and Affiliations

**Department of Human Genetics, Leiden University Medical Center, Leiden, The Netherlands**

Mayke A.C. ten Hoor & Peter Hohenstein

**Division of Nephrology, Department of Pediatrics, Willem-Alexander Children’s Hospital, Leiden University Medical Center, Leiden, The Netherlands**

Mayke A.C. ten Hoor & Jaap Mulder

**Transgenic Facility Leiden, Central Animal Facility, Leiden University Medical Center, Leiden, The**

**Netherlands**

Margot M. Linssen, Conny M. Brouwers, Jill W.C. Claassens & Peter Hohenstein

**Division of Nephrology, Department of Pediatrics, Sophia Children’s Hospital, Erasmus Medical Center, Rotterdam, The Netherlands**

Jaap Mulder

### Contributions

P.H., M.L., C.B., and J.C. designed and generated the mouse models. M.H., P.H., J.M., M.L., conceived and designed the study. M.H. performed the study and analysis. M.H. wrote the manuscript. M.H., P.H., and J.M., revised and edited the manuscript. All authors reviewed the manuscript.

## Ethics declarations

### Ethical approval

The animal care and experimental procedures were approved by the Animal Welfare Body Leiden and are in accordance with the Dutch Experiments on Animals Act and EU Directive 2010/63/EU.

### Supplementary information

Below is the link to the electronic supplementary information. Supplementary file1

