## Supplementary material for "Molecular characterization of Cdh12-SCON conditional knockout mice reveals unexpected splicing changes": sup file 1

**Table S1 | PCR assays with their corresponding primers, Tm and thermocycling program.**

| Assay | Forward primer | Reverse primer | Tm | Program^1^ |
| --- | --- | --- | --- | --- |
| SCON genotyping | GTTTCTCAGCTGCATTCGGAC | TGCAGTCTGTTGGTTAATAGCA | 60°C | 1 |
| CRE genotyping | GCCTGCATTACCGGTCGATGCAACGA | GTGGCAGATGGCGCGGCAACA CCATT | 67°C | 2 |
| Exon2-exon12 | TGGCTGGGTATGGAATCAGT | CGGCTTCTGTGGTGAGAGAG | 67°C | 3 |
| Exon2-SCON | AGCCCCAACAGACTTTAGCC | CTGAGGGAGGGAAAATAACTTCG | 62.2°C | 1 |
| Exon3a-SCON | TGGTGCTGGCACTGTTTTTAC | CTGAGGGAGGGAAAATAACTTCG | 62.2°C | 1 |
| SCON-exon3b | TACGAAGTTATTTTCCCTCCCTC | ACTAGCAACATAAGGTCCATCCA | 62.2°C | 1 |
| SCON-exon4 | TACGAAGTTATTTTCCCTCCCTC | CTTCACCTGCAGCACATACG | 62.2°C | 1 |
| SCON-exon5 | TACGAAGTTATTTTCCCTCCCTC | CCTCCCATGTCTTTCGCTTG | 62.2°C | 1 |
| SCON-exon6 | TACGAAGTTATTTTCCCTCCCTC | ACAGGGGAAGACTCGGGAAC | 62.2°C | 1 |
| SCON-exon7 | TACGAAGTTATTTTCCCTCCCTC | GGTTTGCTGAACACTGGTGG | 62.2°C | 1 |
| Exon2-exon3a | AGCCCCAACAGACTTTAGCC | AGCACGAAGAGTGTAGAAAGGT | 67.5°C | 1 |
| Exon2-exon4 | AGCCCCAACAGACTTTAGCC | CTTCACCTGCAGCACATACG | 67.5°C | 1 |
| Exon2-exon5 | AGCCCCAACAGACTTTAGCC | CCTCCCATGTCTTTCGCTTG | 62.2°C | 1 |
| Exon2-exon6 | AGCCCCAACAGACTTTAGCC | ACAGGGGAAGACTCGGGAAC | 70°C | 1 |
| Exon2-exon7 | AGCCCCAACAGACTTTAGCC | GGTTTGCTGAACACTGGTGG | 70°C | 1 |
| Exon3a-exon4 | TGGTGCTGGCACTGTTTTTAC | CTTCACCTGCAGCACATACG | 70°C | 1 |
| Exon3b-exon4 | GGACATAGAAACCAGGAAGCCA | CTTCACCTGCAGCACATACG | 70°C | 1 |

^1^Program 1 corresponds to 03:00 (95°C); 35 x 00:30 (95°C), 00:30 (Tm), 01:00 (72°C); 05:00 (72°C), program 2 to 05:00 (95°C); 35 x 00:30 (95°C), 00:30 (Tm), 01:00 (72°C); 10:00 (72°C), and program 3 to 03:00 (98°C); 35 x 00:10 (98°C), 00:30 (Tm), 01:20 (72°C); 05:00 (72°C).

**gBlock sequence**

5’CACCACTTCAGCCACAGCCCCAACAGACTTTAGCCACAGAACCAAAAGAAAATGTTATCCACCTTTCGGGGAGACGATCCCATTTCCAACGAGTTAAACGTGGCTGGGTATGGAATCAGTTTTTTGTGCTGGAAGAGTACATGGGCTCCGAACCTCAATATGTGGGGAAGCTGCATTCGGACTTGGATAAAGGAGAGGGCACTGTTAAATACACGCTCTCCGGAGATGGTGCTGGCACTGTTTTTACAATTGATGAAACTACAGGAGACATTCATGCAATAAGAAGCCTGGATAGAGAAGAAAAACCTTTCTACACTCTTCGTGCTCAGGTAAGTAATAACTTCGTATAAGGTATCCTATACGAAGTTATTTTCCCTCCCTCAGGCAGTGGACATAGAAACCAGGAAGCCACTGGAGCCTGAATCAGAGTTCATCATTAAAGTGCAGGATATTAATGACAATGAACCAAAGTTTTTGGATGGACCTTATGTTGCTAGTGTTCCAGAAATGTCTCCTGTGGGTGCGTATGTGCTGCAGGTGAAGGCCACAGATGCAGACGATCCTACCTATGGGAACAGTGCCAGAGTTGTTTACAGCATCCTTCAGGGGCAACCTTATTTCTCTATTGATCCCAAAACAGGTGTCATTAGAACAGCATTGCCAAACATGGACAGAGAAGTCAAAGAGCAGTACCAGGTCCTCATTCAAGCGAAAGACATGGGAGGACAGCTCGGAGGACTTGCTGGAACTACAGTTGTCAACATCACCCTTACTGATGTCAATGACAACCCACCTCGCTTCCCAAAAAGCATCTTCCATCTGAAAGTTCCCGAGTCTTCCCCTGTTGGCTCAGCTATTGGAAGAATAAGAGCAGTAGATCCTGATTTTGGAAAAAATGCAGAAATTGAATACAACATTGTCCCAGGAGATGGGGGAAATTTGTTTGACATTGTCACAGATGAGGATACACAAGAAGGAATCATCAAATTGAAAAAGCCTTTAGATTTTGAAACCAAGAAGGCATATACTTTTAAAGTAGAGGCCTCCAACCTTCACCTTGACCACCGCTTTCACTCTGCTGGGCCATTTAAGGATACTGCTACAGTAAAAATCAGCGTGCTGGATGTGGATGAGCCACCAGTGTTCAGCAAACCACTGTACACCATGGAGGTTTATGAAGACACTCCTGTGGGGACCATCATCGGAGCTGTCACAGCACAAGACCTTGATGTGGGCAGTAGTGCTGTTAG 3’

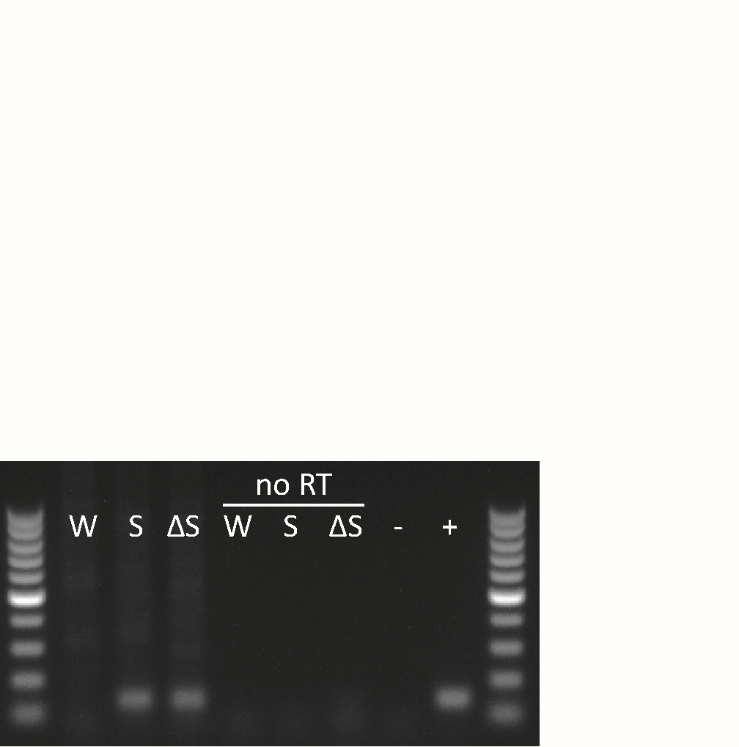

**Figure S1 | gDNA contamination test for (Δ)SCON – exon 3b.** PCR using the (Δ)SCON – exon 3b assay on RNA with and without reverse transcriptase (RT). Marker is 100 bp ladder. W = wildtype, S = Cdh12^SCON/SCON^, ΔS = Cdh12^ΔSCON/ΔSCON^, - = MQ, + = gBlock.
